# Determinants of directionality and efficiency of the ATP synthase F_o_ motor at atomic resolution

**DOI:** 10.1101/2021.09.06.459109

**Authors:** Antoni Marciniak, Pawel Chodnicki, Kazi A Hossain, Joanna Slabonska, Jacek Czub

**Affiliations:** Department of Physical Chemistry, Gdansk University of Technology, Gdansk, Poland; BioTechMed Center, Gdansk University of Technology, Gdansk, Poland

## Abstract

F_o_ subcomplex of ATP synthase is an membrane-embedded rotary motor that converts proton motive force into mechanical energy. Despite a rapid increase in the number of high-resolution structures, the mechanism of tight coupling between proton transport and motion of the rotary c-ring remains elusive. Here, using extensive all-atom free energy simulations, we show how the motor’s directionality naturally arises from the interplay between intra-protein interactions and energetics of protonation of the c-ring. Notably, our calculations reveal that the strictly conserved arginine in the a-subunit (R176) serves as a jack-of-all-trades: it dictates the direction of rotation, controls the protonation state of the proton-release site and separates the two proton-access half-channels. Therefore, arginine is necessary to avoid slippage between the proton flux and the mechanical output and guarantees highly efficient energy conversion. We also provide mechanistic explanations for the reported defective mutations of R176, reconciling the structural information on the F_o_ motor with previous functional and single-molecule data.

---

F_o_F_1_-ATP synthase is a ubiquitous multi-subunit protein that reversibly couples the proton gradient across energy-transducing membranes to the synthesis of ATP from ADP and inorganic phosphate. ^1^ It consists of two mechanically-coupled rotary motors: the hydrophilic F_1_, driven by ATP hydrolysis, and the membrane-embedded F_o_, powered by proton translocation across the membrane. ^2–5^ Two components of F_o_ that are directly involved in this transport are the c-ring, i.e., a rotating oligomer of c-subunits, and the stator a-subunit that wraps around the c-ring (Fig. 1) to form two hydrated proton-access half-channels. ^4,6–10^

During mitochondrial ATP synthesis, protons from the perimitochondrial space bind to the conserved carboxylate near the middle of the c-subunit at proton-binding half-channel (B; yellow in Fig. 1) and are released to the matrix when the same c-subunit reaches the release half-channel (R; red) after almost 360° rotation (magenta pathway). ^9,11–16^ As a result, proton flow through F_o_ induces counterclockwise (when viewed from the matrix) rotation of the c-ring relative to the a-subunit, generating torque that drives the rotation of F_1_ and eventually leads to ATP synthesis. ^17,18^

The question that raises is how F_o_ ensures unidirectional rotation of the c-ring preventing futile proton leakage – a necessary condition for a remarkably high efficiency of the free energy transduction. ^19,20^ Because the rotation by one c-subunit in either direction leads to equivalent states, the synthesis direction has to be kinetically preferred, i.e., the energetic barrier along the mechanical coordinate should be markedly lower in the synthesis direction than in the hydrolysis direction.

**Figure 1.**
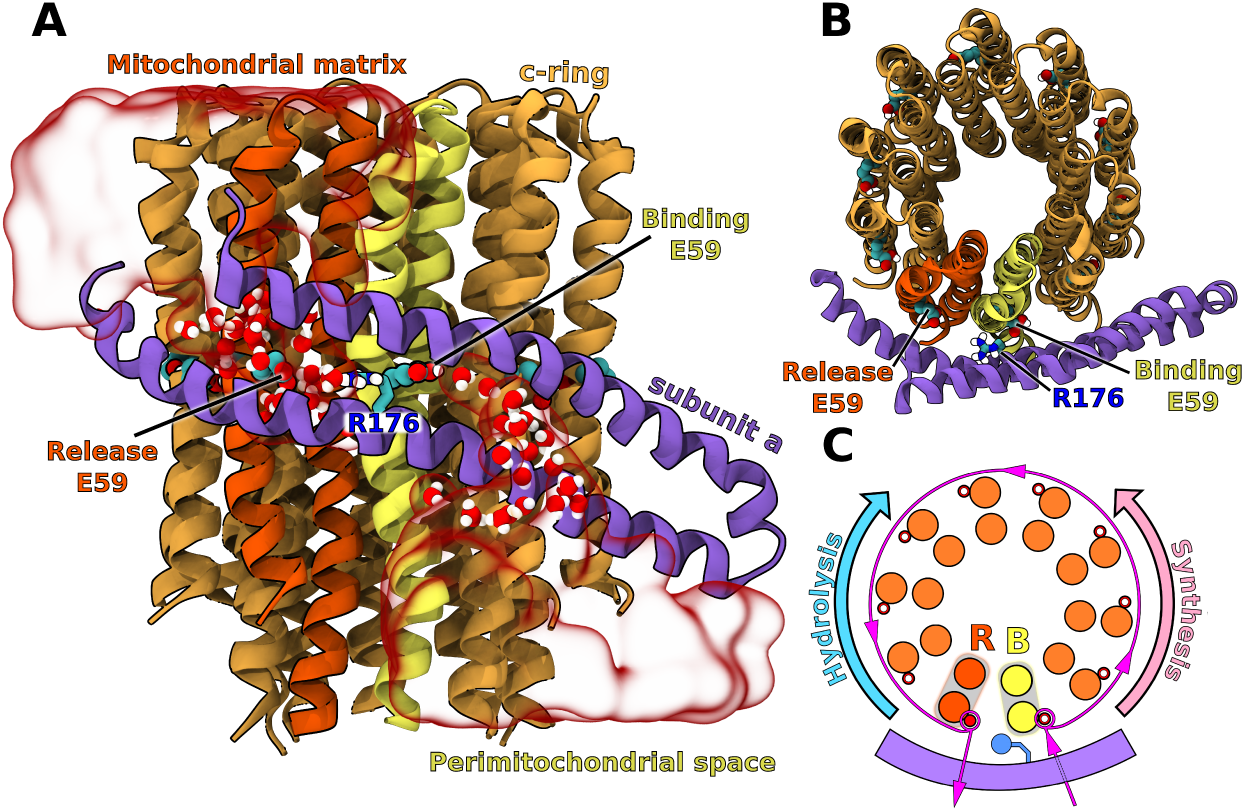
Side (A) and top (B) view of the c-ring/a-subunit interface. Transparent red surface shows representative water distribution in the two half-channels (for average water density, see Fig. S1). The c-subunits shown in red and yellow mark the binding (B) and release (R) sites for protons in the synthesis mode. Only residues 152–249 in the a-subunit are shown for clarity. For full subunit composition of the simulated F_o_ see Fig. S2. (C) Proton transfer pathway through F_o_ in the synthesis mode. In the ATP hydrolysis-driven pumping mode, the c-ring rotates in the reverse direction.

This kinetic preference has been proposed to arise from the specific interactions between the central arginine residue in the a-subunit (R176 in Fig. 1) and carboxylates in the proton binding and release half-channels. ^21,22^ Indeed, mutational studies have shown that the central arginine is essential for coupling proton translocation to mechanical motion and preventing proton leakage. ^23–25^ Because the Arg guanidine moiety is located right in the middle between the R and B half-channels (Fig. 1B), its electrostatic attraction to the freshly deprotonated carboxylate at site R should favor the rotation in the synthesis direction and oppose the reverse rotation. ^26–28^ By determining the free energy landscapes using a coarse-grained model, Bai and Warshel have recently shown that this simple electrostatic view of the F_o_ directionality only holds if the energetics of proton transfer to the carboxylates is included, ensuring the asymmetric attraction between Arg and the R and B sites. ^29,30^ While these findings explain the general aspects of the mechanochemical coupling, they do not provide atomic-level understanding of the F_o_ mechanism; in particular, it is not known why Arg is strictly conserved and cannot be substituted by a positively charged Lys residue that could act in an analogous manner. Here, by using atomistic molecular dynamics-based free energy simulations (total sampling time of nearly 70 *µ*s), we provide a detailed insight into the origin of the directionality of the c-ring rotation and elucidate the role of the central arginine in the F_o_ mechanism.

To this end, we determined and compared the free energy profiles for the rotation of the yeast mitochondrial F_o_ by one c-subunit in the synthesis and hydrolysis directions (+36° and − 36°, respectively in Fig. 2; for details see Supporting Information (SI) Methods). Because energetics of the c-ring rotation has been found to strongly depend on the protonation state of the B and R carboxylates, we first calculated their pKa’s in the F_o_ initial state (0°) using the alchemical approach (SI Methods). We found (inset in Fig. 2) that pKa of the B site (8.4±0.3) is markedly higher and that of the R site (1.9±0.2) is markedly lower compared to pH of their respective compartments (e.g., 6 and 7, respectively), even if the predicted shifts are somewhat overestimated. This finding suggests that proton affinities for the B and R sites are fine-tuned to the proton gradient direction, promoting the rate of the proton transfer via F_o_ under physiological conditions. As can be seen, the free energy profiles determined for the dominant protonation state (deprotonated R, protonated B, R^−^ B^0^; blue curves in Fig. 2) are locally strongly asymmetric around 0°, implying a ∼15 kcal/mol kinetic preference for the rotation in the synthesis direction, consistently with the motor’s unidirectional motion.

**Figure 2.**
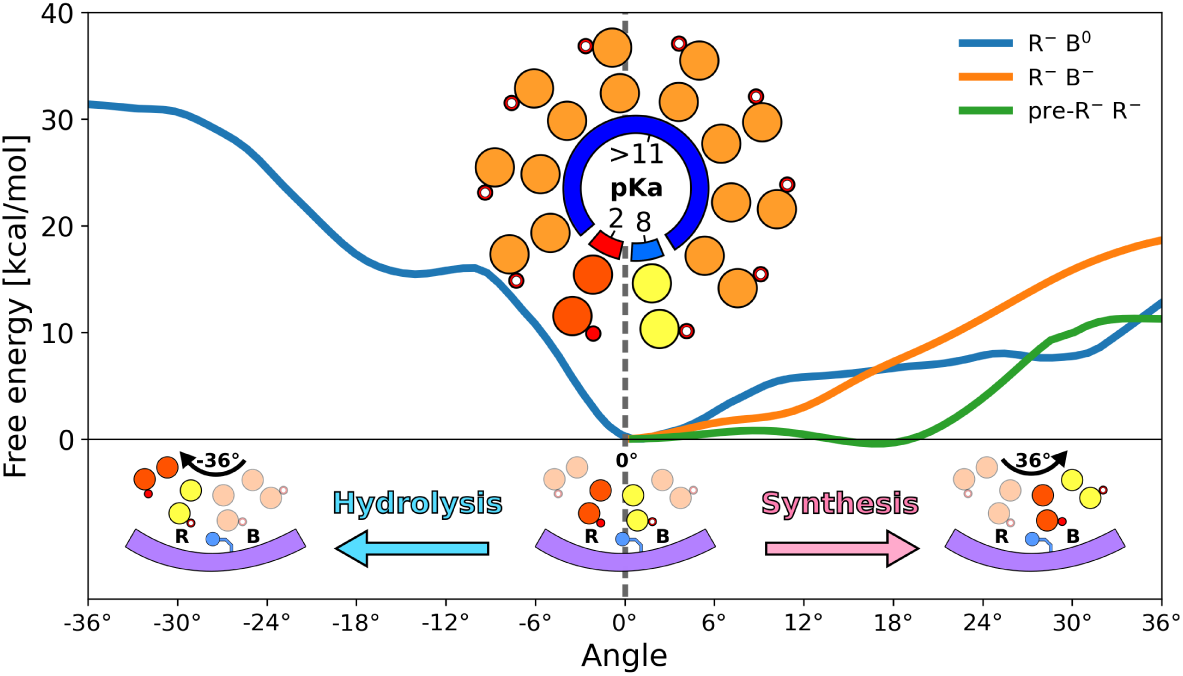
Free energy profiles for the F_o_ rotation in hydrolysis and synthesis direction by one c-subunit (0 → 36°), at different protonation states of the c-ring. Inset in the middle shows the computed pKa’s of the c-ring carboxylates (at 0° position, as captured by the cryo-em structure). ^5^ For convergence of the free energy profiles, see Fig. S3–S5

It has been suggested, however, that not only protonation thermodynamics (as described by pKa’s) but also the rate of (de)protonation might play an important role in the directional rotary mechanism. ^30^ In particular, protonation of the B site could even be expected to be a rate limiting step, given that a single file of water molecules confined between the helices of the a-subunit connects this site to the rest of the half-channel (as seen in Fig. 1 and Fig. S1, consistently with previous cryo-EM data). ^7,31,32^ Therefore, we examined the effect of the B site deprotonation on the rotation energetics, by recomputing the free energy of the rotation in the synthesis direction with both R and B sites deprotonated (R^−^ B^−^; orange curve in Fig. 2).

As can be seen, beyond 10° the R^−^ B^−^ profile rapidly increases with a roughly constant slope of 1.8 kcal/(mol · deg), reflecting the change of environment around the B site carboxylate from polar and partially hydrated (at 0°) to fully hydrophobic (at 36°). This increase effectively inhibits the full rotary step in this protonation state ensuring that the c-ring waits, if necessary, for the protonation of the B site to occur. Surprisingly, however, in the range up to 10°, the R^−^ B^−^ protonation state seems to kinetically favor the synthesis direction even more than the R^−^ B^0^ protonation state, as the free energy barrier for the former is ~3 kcal/mol lower. Since the two curves change their slope around 10°, with R^−^ B^−^ becoming notably steeper and R^−^ R^0^ flatter, it may suggest that the c-ring, starting from 0°, first tends to rotate with the B site carboxylate deprotonated up to 10°, where it stalls waiting for the carboxylate to get protonated, to progress further governed by the less steep R^−^ B^0^ profile. This hypothesis would agree with the recent single-molecule study that revealed the existence of a 11° substep in the rotary mechanism of the *E*.*coli* F_o_. ^33^ A doubly deprotonated state of the c-ring was also found to contribute to the proton transfer pathway in the recent coarse-grained simulations. ^13^

Because during the rotation in the synthesis direction the c-subunit entering the R site (pre-R subunit) moves from hydrophobic to polar and well-hydrated environment (Fig. S1 and Fig. S6), which corresponds to a drastic decrease in the pKa of E59, early deprotonation of the pre-R carboxylate can also be expected to affect the rotation kinetics. To test this possibility, we determined the effect of deprotonation of the pre-R carboxylate on the synthesis free energy profile [(pre-R)^−^ R^−^; green curve in Fig. 2]. We found that for the (pre-R)^−^ R^−^ state, as opposed to the two other examined protonation states, the profile remains virtually flat up to 20°, which shows that early proton release might in fact promote faster progression of the c-ring in the synthesis direction, further highlighting the coupling between rotation and (de)protonation events.

Next, to gain atomic-level understanding of the directionality mechanism, we examined residue-residue and residue-environment enthaplic contributions to the interaction free energies along the mechanical coordinate (see SI Methods for details). In Fig. 3, the average local slopes of the corresponding contributions in the synthesis and hydrolysis direction were subtracted from each other, such that the resulting negative values indicate the interactions favoring the c-ring rotation in the synthesis direction over the hydrolysis direction, and vice versa. It can be clearly seen that the attractive interaction between the deprotonated E59 at the release site (R^−^) and the central R176 is a key differentiating factor largely responsible for the directional preference in the dominant R^−^ B^0^ protonation state, supporting the electrostatic model of the F_o_ unidirectionality. ^30^ Indeed, the intra-protein interactions involving R^−^ and R176 are 4.25 and 3 times, respectively, more effective in promoting the synthesis direction than those involving the 3rd most contributing residue, i.e., also highly conserved R169, whose interaction with R^−^ seem to favor the synthesis too. Since strengthening of the electrostatic attraction between pairs of oppositely-charged residues over the course of the rotation is accompanied by their dehydration, the interactions of R^−^, R176 and R169 with water partially counteract the intra-protein preference by favoring the hydrolysis direction (see Fig. S10). In all cases, interactions with lipid molecules have mostly negligible effect on directionality.

**Figure 3.**
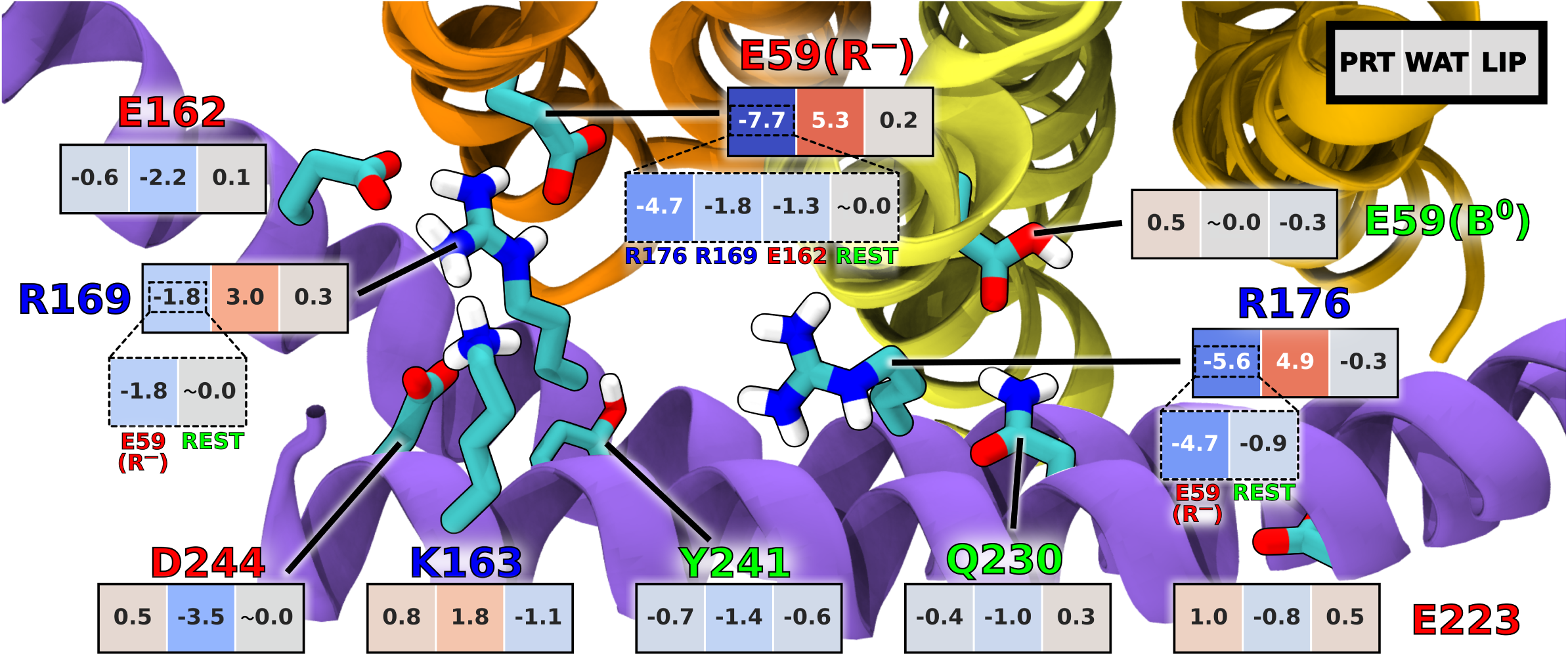
Comparison of the per-residue enthalpic contributions to the rotation free energies in the synthesis and hydrolysis direction. Values shown are the differences in the average slopes of these contributions with respect to the rotation angle between the synthesis and hydrolysis directions, calculated in the 0–18° range. Accordingly, negative values indicate the contributions favoring the rotation in the synthesis direction and vice versa. Interactions of each residue within the protein (PRT), with water (WAT) and lipids (LIP) are shown separately. Only residues with any of the slope differences exceeding 1.0 kcal/(mol·deg) are included (with exception of E59(B^0^); see Fig. S7–S10 for complete pairwise data).

To directly test the pivotal role of the central arginine (here R176) in differentiating the rotation energetics in the synthesis vs. hydrolysis direction, we further determined how its mutation to alanine (R176A) affects the free energy landscape governing the rotary motion of the c-ring around 0° (Fig. 4A). As seen from the comparison with the wild type profiles, the R176A mutation leads to a striking reduction of the barrier to rotation in the hydrolysis direction (by ∼15 kcal/mol), while having only a limited effect on the free energy in the synthesis direction. Thus, upon removal of the central arginine, the energy landscape becomes roughly symmetric around 0° implying almost no kinetic preference for either of the directions (in fact, hydrolysis seems to be slightly favored). Similar rotation rates in both directions indicate that the R176A mutant should show a severe slippage between the proton flux and the mechanical output (and, consequently, ATP synthesis), as a substantial fraction of protons would be transported to the R site and released after only 36° rotation in the hydrolysis direction. This finding provides a possible mechanistic explanation for the rotation-uncoupled proton transport observed previously upon this mutation ^24,34^ and could be directly tested in single-molecule imaging experiments.

**Figure 4.**
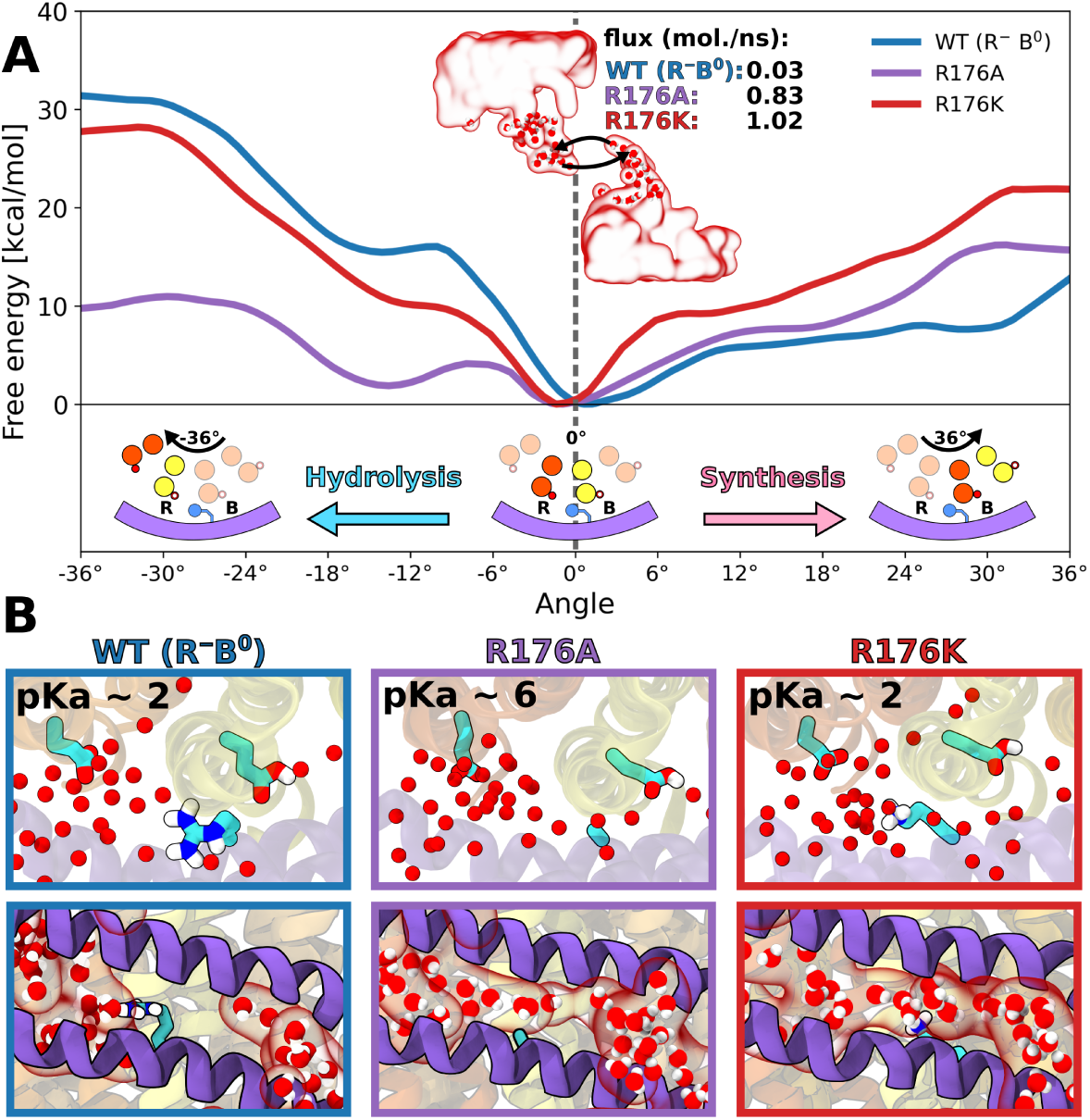
(A) Response of the free energy profile for the F_o_ rotation to the mutations of the central arginine (R176). Inset shows the calculated water flux between the two half-channels. (B) (top) calculated pKa’s of the R-site carboxylate and (bottom) representative distribution of water molecules around the position 176, for the WT protein and the two arginine substitutions. For convergence of the free energy profiles, see Fig. S3 and Fig. S11–S12

Comparison of the residue-residue interactions between the wild type and R176A mutant (see Fig. S8) further substantiates the critical role of electrostatic attraction between the central arginine and the deprotonated R-site carboxylate in promoting the unidirectional rotation in the synthesis direction. This simple mechanism implies that any pair of oppositely charged residues could work in a similar manner. Indeed, the c-ring glutamate (here E59) can occasionally be replaced by aspartate without abolishing the directionality, e.g., in some Gram-negative bacteria, including *E*.*coli*. In contrast, arginine cannot be replaced by any other residue, including lysine, i.e., the only other residue positively charged at physiological conditions. To explore why this substitution might be detrimental, we first determined and compared the hydrolysis and synthesis free energy profiles for the lysine mutation (R176K) that was also previously shown to inhibit the ATP-driven proton pumping activity. ^24^ It is clear from Fig. 4A that the synthesis profile is markedly steeper for R176K than for the wild type protein, such that the barrier to rotation in the synthesis direction is much more pronounced; e.g., for the 0–6° range the barrier is ∼7.5 kcal/mol higher which renders the rotation rate five orders of magnitude slower compared to WT. Since electrostatic attraction between K176 and the R-site carboxylate (see Fig. S9)) preserves a relatively high (∼12 kcal/mol) barrier to the rotation in the hydrolysis direction (Fig. 4A), we predict that in the R176K mutant the c-ring cannot rotate away from 0° in either of the directions on physiologically relevant time scales. This finding explains the previous reports that the R176K mutant is unable to couple the proton motive force to ATP synthesis but, at the same time, it does not exhibit futile proton leakage as that caused by the R176A mutation. ^24^

Because the side chain of K176 is much more flexible than that of R176 (root mean square fluctuation of 0.072 vs. 0.021 nm^2^, respectively), it could be expected to interact with the carboxylate in the release site less favorably, and thus lead to an undesired increase of its pKa. To test this, we computed pKa of the R-site carboxylate in the R176K mutant and, as a control, in the R176A mutant (Fig. 4B). We found that both positively charged residues, arginine and lysine, stabilize the deprotonated state of the carboxylate to a similar extent, lowering its pKa by ∼4 units from a value of 6.9±0.1 calculated for the R176A mutant to 1.9±0.2 (WT) and 2.2±0.4 (R176K). Since the effect on pKa of the R-site carboxylate does not depend on the exact nature of a positively charged residue, we conclude that it alone cannot explain strict conservation of the central arginine.

Having found no significant differences in the interactions with the R-site carboxylate, we turned to examining whether arginine and lysine, given their position in between the B and R site, may cause different behavior of water molecules in the two half-channels. To this end, we determined average water densities in the F_o_ half-channels for the wild type protein as well as for both mutants: R176K and R176A. As can be seen in Fig. 4B and Fig. S1, the arginine side chain at position 176 acts as a “plug” that separates water density into two well-defined half-channels. In contrast, in both mutants the half-channels are not disconnected anymore but form a continuous aqueous pore through which water molecules can rapidly diffuse across the membrane. Specifically, we calculated the water flux between the two half-channels to be two orders of magnitude larger in both mutants than in the wild type (1.02, 0.83 and 0.03 molecules/ns, for R176K and R176A and WT, respectively). One could expect that the connected channel can also allow protons to leak across the membrane via Grotthuss mechanism downhill the gradient in a manner uncoupled from the c-ring rotation. Such rotation-uncoupled proton leakage has indeed been observed for the alanine mutant by Mitome et al., who showed that, even when the c-ring is fused with the a-subunit and thus unable to rotate, the F_o_ can still act as a proton channel when the central arginine is substituted by alanine. ^24^ Although the continuous water channel is also predicted to be present in the R176K mutant (Fig. 4B, bottom), the lysine mutation has not been observed to lead to the uncoupled proton leakage. ^24^ We hypothesize that in this case protons are unable to move directly through the continuous channel from the binding to release site because of the electrostatic repulsion with the lysine.

To sum up, we found that the free energy landscape governing the F_o_ rotation is highly asymmetric around the initial position of the c-ring (0°), and thus rotation in the synthesis direction is strongly preferred, consistently with the motor’s directionality and high efficiency of energy conversion. Importantly, this kinetic preference arises from the interplay between the intra-protein interactions and the energetics of protonation of carboxylates in the binding (B) and release (R) half-channels. Specifically, we predicted that at 0° the B site is predominantly protonated and the R site deprotonated, and hence their pKa’s seem to be fine-tuned to the direction of the proton gradient, accelerating the proton flow through F_o_ under physiological conditions. Our energetic analysis demonstrated that in this dominant protonation state the main contribution accounting for the much lower activation barrier in the synthesis direction is the attraction between the strictly conserved arginine in the asubunit (R176 in the yeast F_o_) and the deprotonated carboxylate at the R site. By recomputing the free energies for the R176A mutant, we further confirmed this finding, showing that in the absence of R176 the directionality of F_o_ is largely abolished and the pKa of the R site unfavorably increases by 4 units. Furthermore, the alanine mutation causes the B and R aqueous half-channels, that are well-separated by R176 in the wild type, to join in a single continuous pore through which water can readily move across the membrane. Along with the abolished directionality, the existence of this shortcut provides mechanistic explanation for the previously reported uncoupled proton transport caused by removal of the arginine side chain. Finally, we found that a positively charged residue, lysine, substituted for R176 facilitates deprotonation of the R site and preserves a high barrier to rotation in the hydrolysis direction similarly to the arginine. However, it also markedly increases the barrier in the synthesis direction which slows down the rotation by several orders of magnitude, further explaining why arginine has to be conserved to ensure efficient coupling between proton translocation and rotary motion in the F_o_ motor.

## Acknowledgement

We thank Dr. Mi losz Wieczór for helpful discussions and critically reading the manuscript. This work was funded by the Polish National Science Centre under Sonata Bis grant No. 2017/26/E/NZ2/00472. This research was supported in part by PL-Grid Infrastructure. Computational resources were provided also by the TASK, WCSS and ICM Centers.

