## Supplemental Information for "Determinants of directionality and efficiency of the ATP synthase F_o_ motor at atomic resolution"

### Supporting Information for: Determinants of directionality and efficiency of the ATP synthase $F_o$ motor at atomic resolution

Antoni Marciniak,† Pawel Chodnicki,† Kazi A Hossain,† Joanna Slabonska,† and Jacek Czub*,†,‡

†*Department of Physical Chemistry, Gdansk University of Technology, Gdansk, Poland*

‡*BioTechMed Center, Gdansk University of Technology, Gdansk, Poland*

#### Methods

##### System preparation

The cryo-EM structure (PDB Id: 6B2Z)[1] of yeast mitochondrial ATP synthase (isolated $F_o$ monomer) was taken as an initial configuration. As e- and g-subunit, located far from the crucial a/c-ring interface, were modeled in the cryo-EM structure as poly-Ala chains, they were completely omitted. Using the CHARMM-GUI Membrane Builder,[2–4] the protein was embedded in a lipid bilayer oriented perpendicularly with respect to the $z$-axis and composed of 151 POPC, 118 POPE, and 55 $TOCL^{-2}$ (tetraoleoyl cardiolipin) molecules (46.6, 36.4 and 17 mol %, respectively), reflecting the composition of the inner mitochondrial membrane in yeast.[5] The system was solvated with 25201 TIP3P water molecules in a $123.85 \times 123.85 \times 91.422$ Å rectangular box and the number of $K^+$ and $Cl^-$ ions was adjusted to maintain a physiological salt concentration of 0.15 M and neutralize the net charge of the system. One POPC and one POPE molecule per leaflet were manually inserted inside the c-ring, to seal its central channel, similarly to the previous treatment.[6] System was subjected to two-step minimization (first keeping all protein atoms fixed and then without any constraints) followed by preliminary relaxation of the protein using CHARMM-GUI protocol (short MD runs with progressively weaker restraints on protein atoms). Next, the system was simulated with the protein backbone atoms harmonically restrained to their initial positions with a force-constant of $1000\,\mathrm{kJ/(mol{\cdot}nm^2)}$ for 500 ns, and only then equilibrated without restraints for another 700 ns. Random frames from the last 100 ns of this restraint-free run were picked as initial coordinates for five independent unbiased MD simulations of the wild type $F_o$, each lasting 1 $\mu$s. Two mutants of $F_o$, R176A and R176K, were prepared by substituting R176 residue with alanine and lysine, respectively, and subject to 2 $\mu$s of unbiased MD simulations, using random frames from the WT equilibrium trajectory as initial coordinates. In the systems where changes in protonation states or mutations created non-zero net charge, the number of ions in aqueous phase was adjusted to neutralize the system.

#### Simulation parameters

All molecular dynamics (MD) simulations were performed using GROMACS[7] and the CHARMM36m force field.[8] The simulations were carried out in the isothermal-isobaric (NPT) ensemble using periodic boundary conditions in 3D. The constant temperature was kept at 310 K using the Nose-Hoover thermostat[9] and the pressure was maintained at 1 bar semi-isotropically (separately in the plane of the bilayer and perpendicular to the bilayer) using Parrinello-Rahman algorithm.[10] Long-range electrostatic interactions were evaluated using the Particle Mesh Ewald (PME)[11] method with a real-space cut-off of 1.2 nm. Van der Waals interactions were evaluated using a smooth cut-off of 1.2 nm with a switching distance of 1 nm. Bond lengths were constrained using the SHAKE[12] (for water) or P-LINCS[13] (for protein and lipids) algorithm. The equations of motion were integrated using a leap-frog algorithm with a time step of 2 fs.

#### $pK_a$ calculations

To determine pKa values of the carboxylates at the binding and release sites of the c-ring in the wild-type and mutant $F_o$, we used alchemical free energy calculations. To this end, a free glutamic acid molecule capped with N-methyl (NME) and acetyl (ACE) termini was added to the $F_o$ system as a reference, and kept in bulk solution, at a minimal distance of 3 nm from the membrane center with the use of a one-sided harmonic potential applied to the $z$ component of the vector connecting the centers of mass of the reference glutmate and the c-ring using PLUMED (topologies were generated using our in-house script (`https://gitlab.com/KomBioMol/proton_alchemist`).[14] To predict the pKa shift for a given c-ring carboxylate with respect to the reference glutamate, the system was transformed, using a switching parameter $\lambda$, from the initial state with the c-ring carboxylate protonated and the reference glutamate deprotonated (state A) to the final state with the reversed protonation states (state B), and vice versa. Due to this treatment the simulated system was neutral at all $\lambda$-points and the obtained $\Delta\Delta$G values, describing the difference in the proton affinity between both environments, could be used directly to compute the pKa shift with respect to the experimental pKa in aqueous solution (4.1 for the glutamic acid side chain). The number of $\lambda$-points (or windows) was chosen to be 20 and the system was simulated in these windows using Hamiltonian replica-exchange molecular dynamics until the convergence of $\Delta\Delta$G was reached ($\sim$200 ns). To optimize $\lambda$ values, we used our in-house script (`https://gitlab.com/KomBioMol/converge_lambdas`) that iteratively restarts short replica-exchange runs until $\lambda$-values yielding equal exchange rates between neighboring windows (here $\sim$20%)) are found.[15] $\Delta\Delta$G values were determined using Bennett acceptance ratio,[16] as implemented in the Gromacs package and the $pK_a$ shifts were then calculated according to the formula $\Delta pK_a = \log_{10}\left(\exp^{\frac{-\Delta\Delta G}{RT}}\right)$

#### Free energy profiles for the c-ring rotation

To determine the changes in the free energy accompanying the rotation of the c-ring by one c-subunit in the wild-type $F_o$ and its R176A and R176K mutants, the umbrella sampling (US) technique was applied. As a reaction coordinate, $\theta$, we used the angle of rotation of the c-ring about the fixed vertical axis passing through its center. For each protein, to span the range of $\theta$ corresponding to the rotation by c-subunit (0–36° and −36–0° for the synthesis and hydrolysis direction, respectively) we used 13 equally-spaced US 'windows' separated by 3°. The stationary isotropic potential, as described by Kutzner et al.,[17] with a force constant of 0.5 kJ/(mol·nm$^2$) was used to restrain the systems in each of these windows. The initial configurations for the US windows were taken from the enforced-rotation simulations in which the c-ring was driven to rotate at a constant angular rate of 0.36 deg/ns by an externally applied isotropic potential with a force constant of 500 kJ/(mol·nm$^2$). To mimic the effect of the peripheral stalk connecting the $F_o$ and $F_1$ portion and prevent the $F_o$ protein to rotate as a whole, the backbone atoms of the membrane-embedded portion of b-subunit (see Fig. S2) were harmonically restrained to their initial positions with a force-constant of 1000 kJ/(mol·nm$^2$). Each of the US windows were simulated for 600 ns (with the exception of the R176K mutant, which was run for 700 and 800 ns in the synthesis and hydrolysis direction, respectively) and first 200 ns were omitted from analysis. Free energy profiles were determined using the multistate Bennett acceptance ratio (MBAR) method, as implemented in pymbar.[18]

#### Interaction analysis

To evaluate the individual enthalpic contributions to the rotation free energies, we calculated electrostatic and van der Waals (vdW) interaction energies between the key residues present at the interface between the c-ring and a-subunit, the remaining protein residues, solvent including ions, and lipids. The binding and release site glutamates and the a-subunit residues within 1 nm from the R site carboxylate forming the interface were selected for the residuewise analysis (for a complete list of residues and their location at the c-ring/a interface, see Fig. S13). Using Boltzmann-reweighted umbrella sampling data, we computed the average electrostatic and vdW interaction energies between all the pairs considered as a function of the rotation angle. After adding electrostatic and vdW energies, the average slope of the resulting angle-dependence in the 0–18° range was determined by linear fitting, and was used as measure of a given pairwise contribution (Fig. S7, Fig. S8, Fig. S9 and Fig. S10). To extract contributions responsible for the $F_o$ directionality the slopes in the hydrolysis direction were subtracted from those in the synthesis direction and shown in Fig. 3.

Table S1: Range of residues included in the simulation for each of the simulated subunits of the $F_o$ complex

| Subunit | range |
|---|---|
| 8 | 8–48 |
| a | 27–249 |
| d | 240–283 |
| f | 32–86 |
| j | 1–36 |
| k | 5–28 |
| b | 8–103 |
| c (×10) | 1–75 |

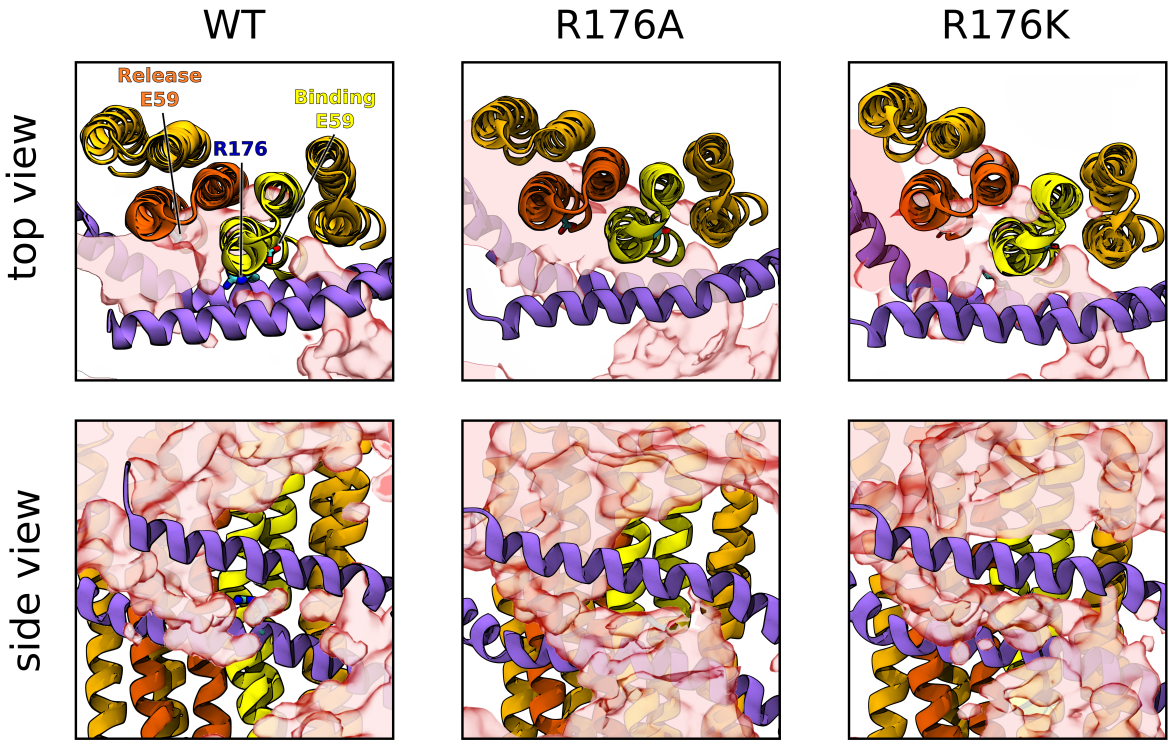


Figure S1: Top and side view of the average water densities (transparent red surfaces) at the interface between a-subunit and the c-ring in the wild type $F_o$ protein, as well as in its R176A and R176K mutants.

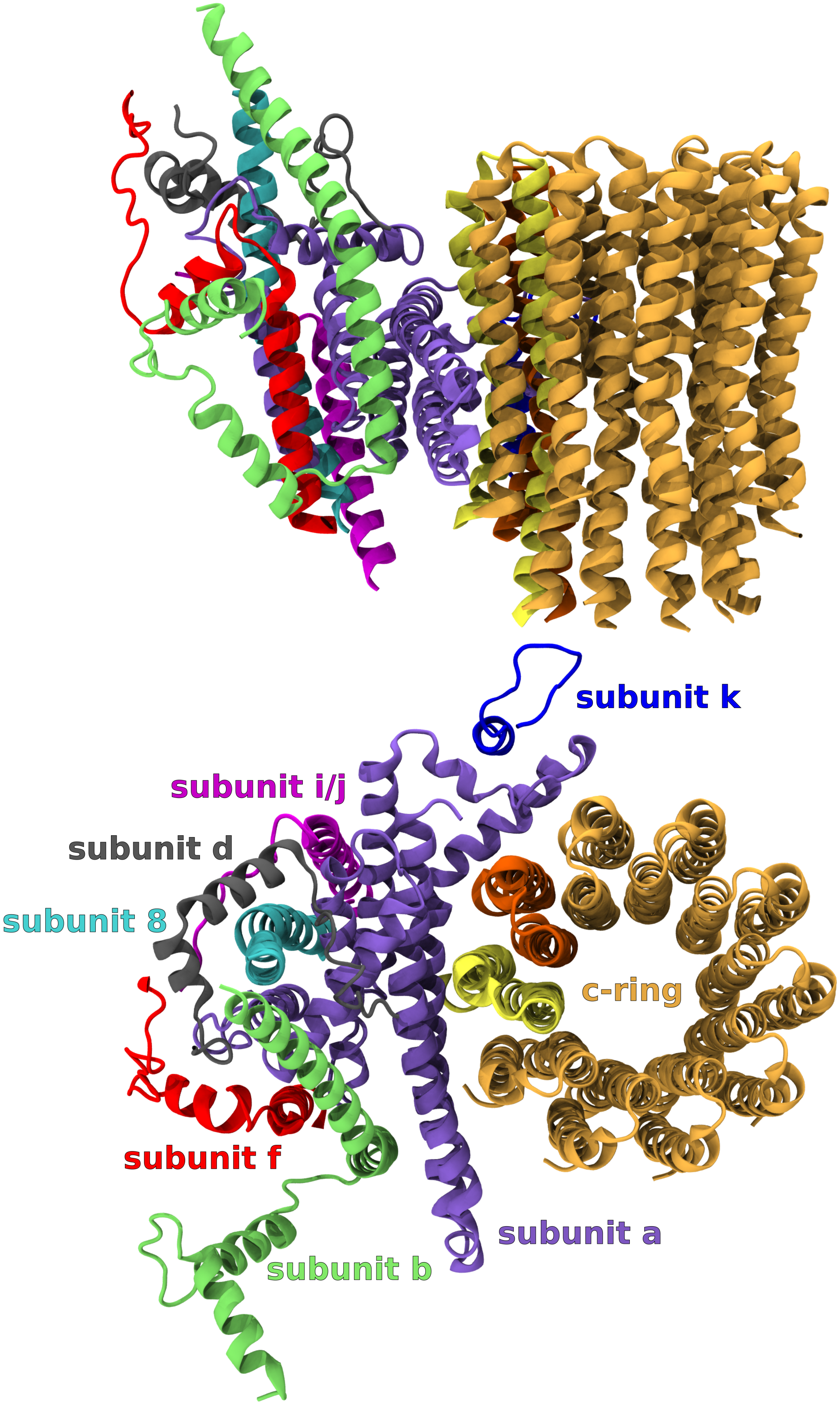


Figure S2: Subunit composition of the simulated $F_o$ complex (top – side view, bottom – view from the mitochondrial matrix). Consistently with Fig. 1., the two c-subunits located in the proton-binding and proton-release half-channels are shown in yellow and red, respectively.

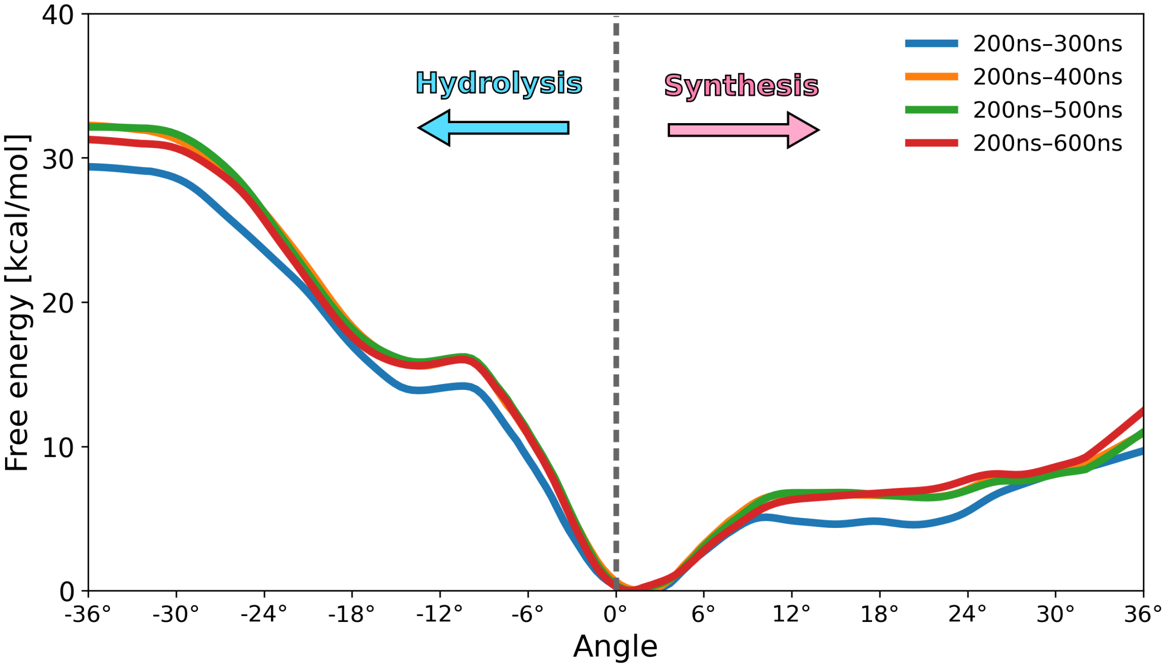


Figure S3: Convergence of the free energy profiles for the rotation of the wild type $F_o$ ($R^-B^0$) in the synthesis and hydrolysis direction, with the length of umbrella sampling trajectories taken for analysis.

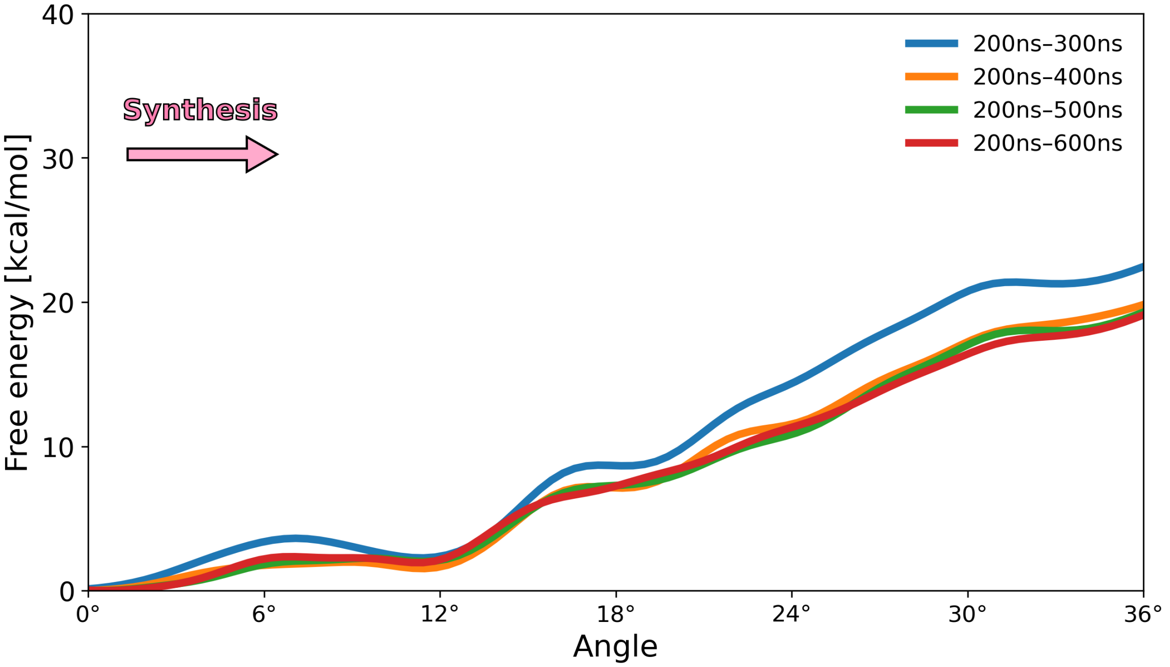


Figure S4: Convergence of the free energy profile for the rotation of the wild type $F_o$ ($R^-B^-$) in the synthesis direction, with the length of umbrella sampling trajectories taken for analysis.

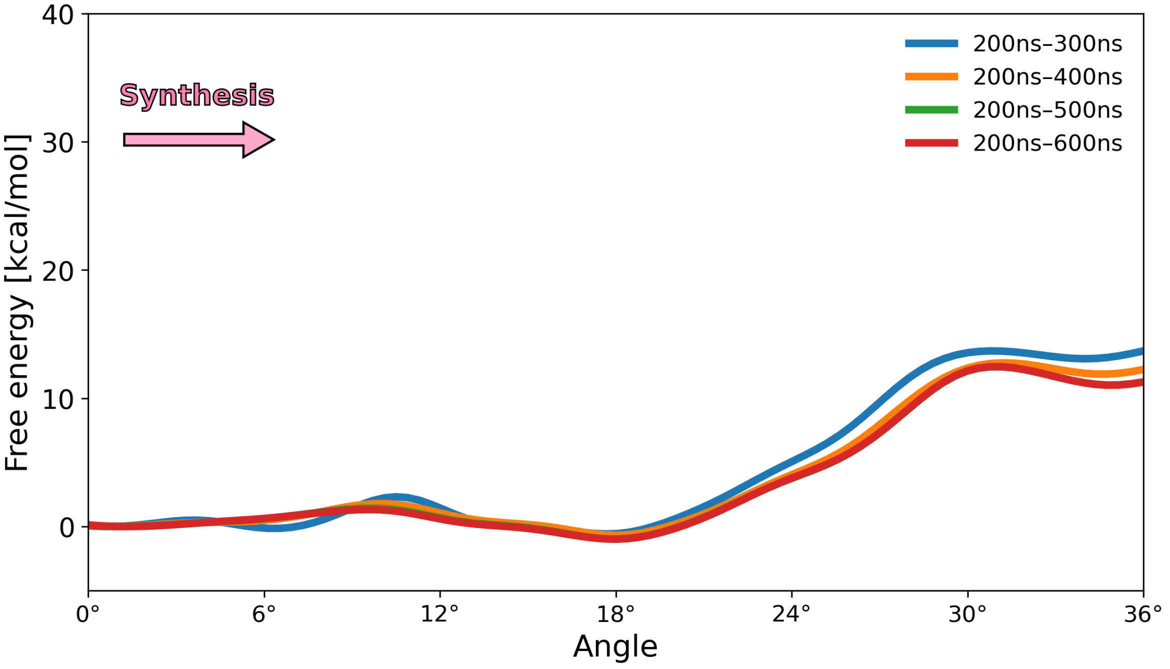


Figure S5: Convergence of the free energy profile for the rotation of the wild type $F_o$ ((pre-R)$^-$R$^-$) in the synthesis direction, with the length of umbrella sampling trajectories taken for analysis.

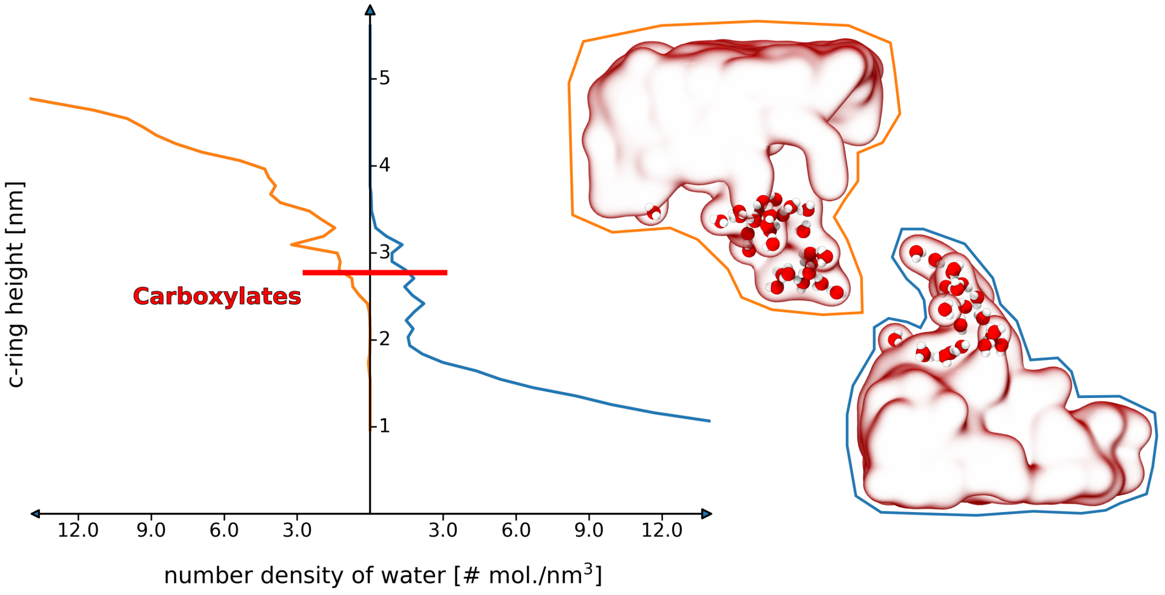


Figure S6: Number density of water in the proton-access half-channels as a function of the c-ring height. Density in the proton-binding and proton-release half-channel are shown in blue and orange, respectively. Red line indicates the height at which the c-ring carboxylates (B and R) are located.

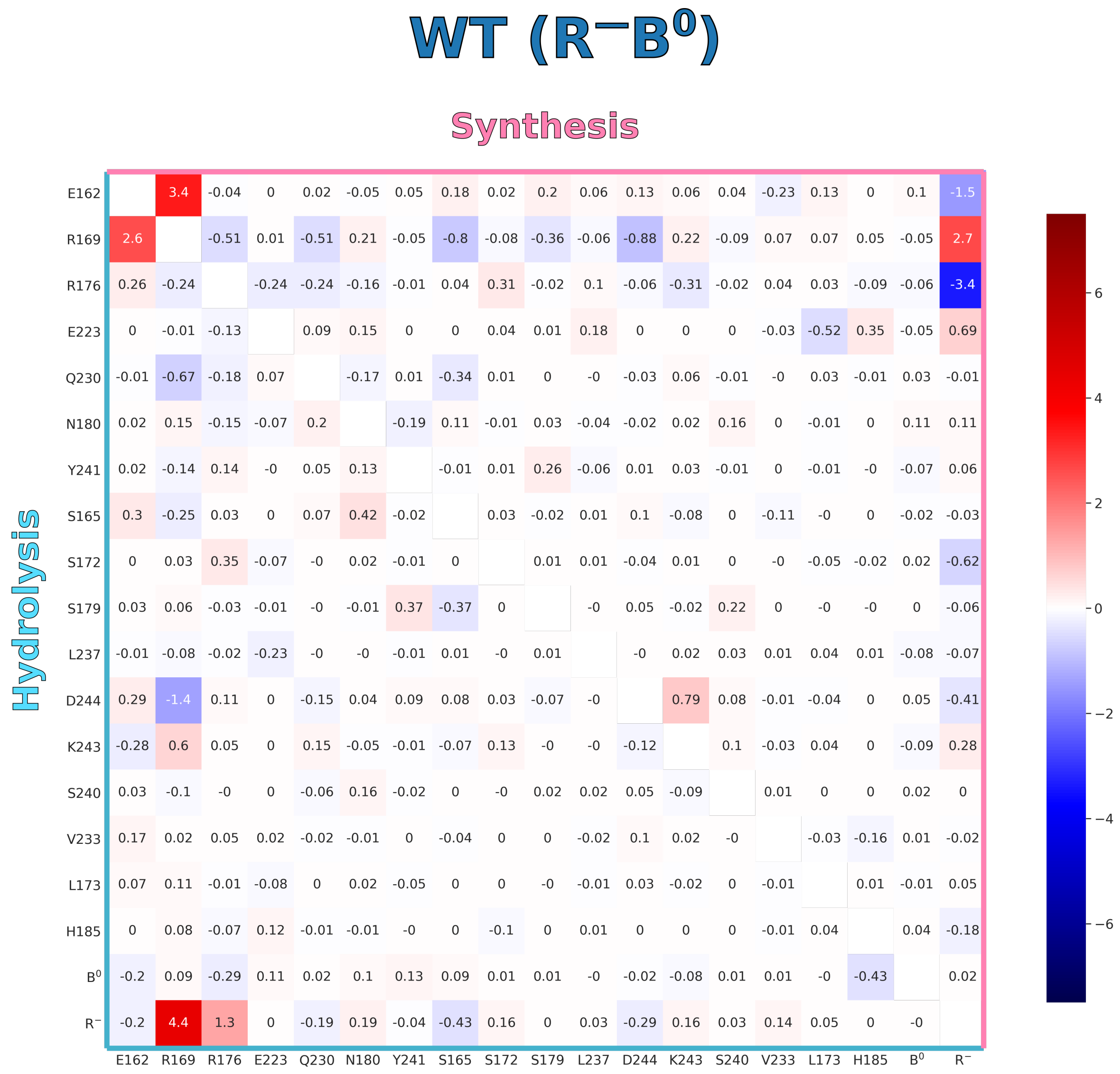


Figure S7: Average slopes (in kcal/(mol·deg)) of the selected pairwise interactions (electrostatic + vdW) as a function of the rotation angle, calculated in the 0–18° range. The values shown in the upper and lower triangle correspond to the rotation of the wild type $F_o$ ($R^-B^0$) in the synthesis and hydrolysis direction, respectively. Locations of the residues used in the analysis are shown in Fig. S13.

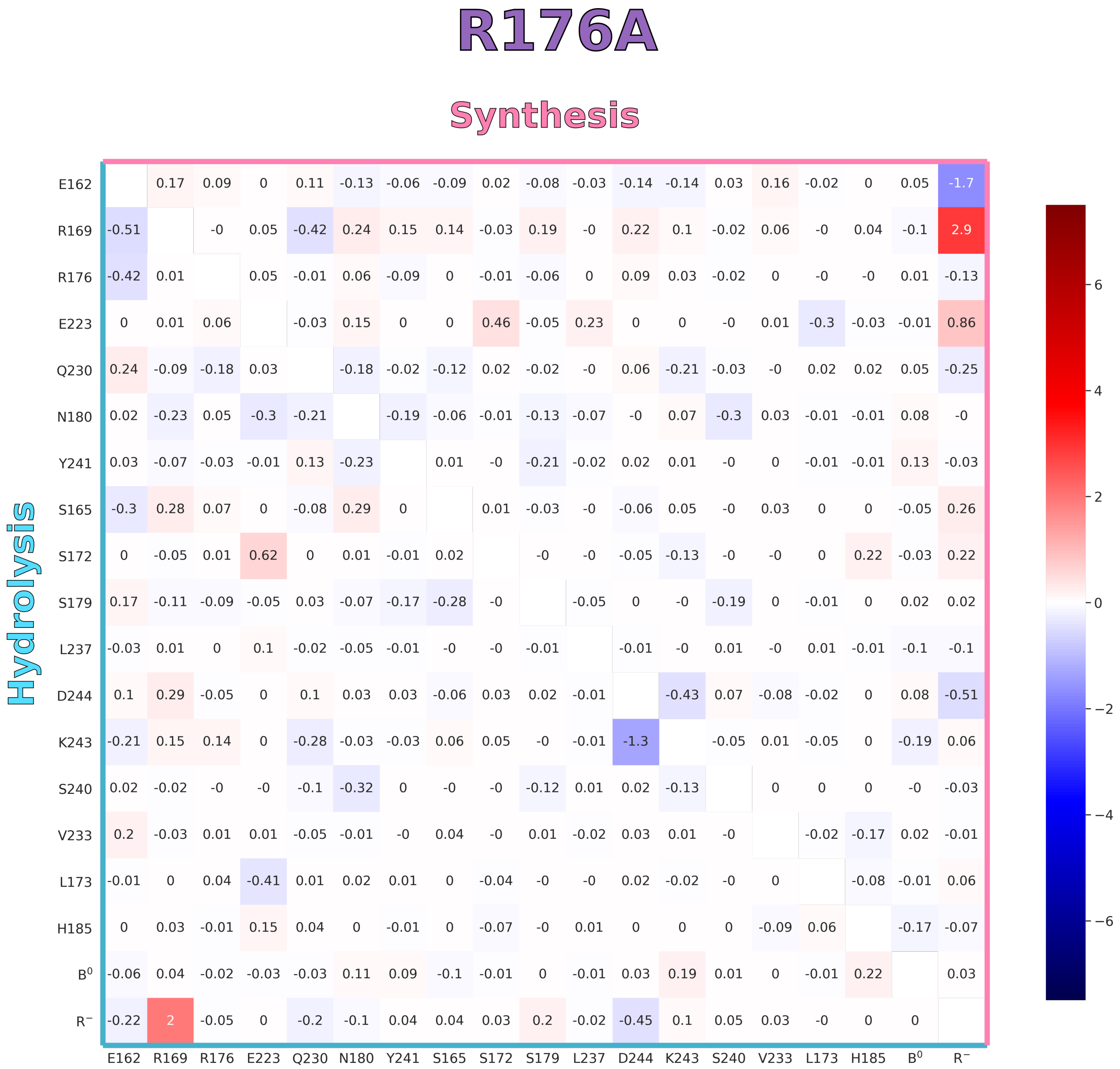


Figure S8: Average slopes (in kcal/(mol·deg)) of the selected pairwise interactions (electrostatic + vdW) as a function of the rotation angle, calculated in the 0–18° range. The values shown in the upper and lower triangle correspond to the rotation of the $F_o$ R176A mutant in the synthesis and hydrolysis direction, respectively. Locations of the residues used in the analysis are shown in Fig. S13.

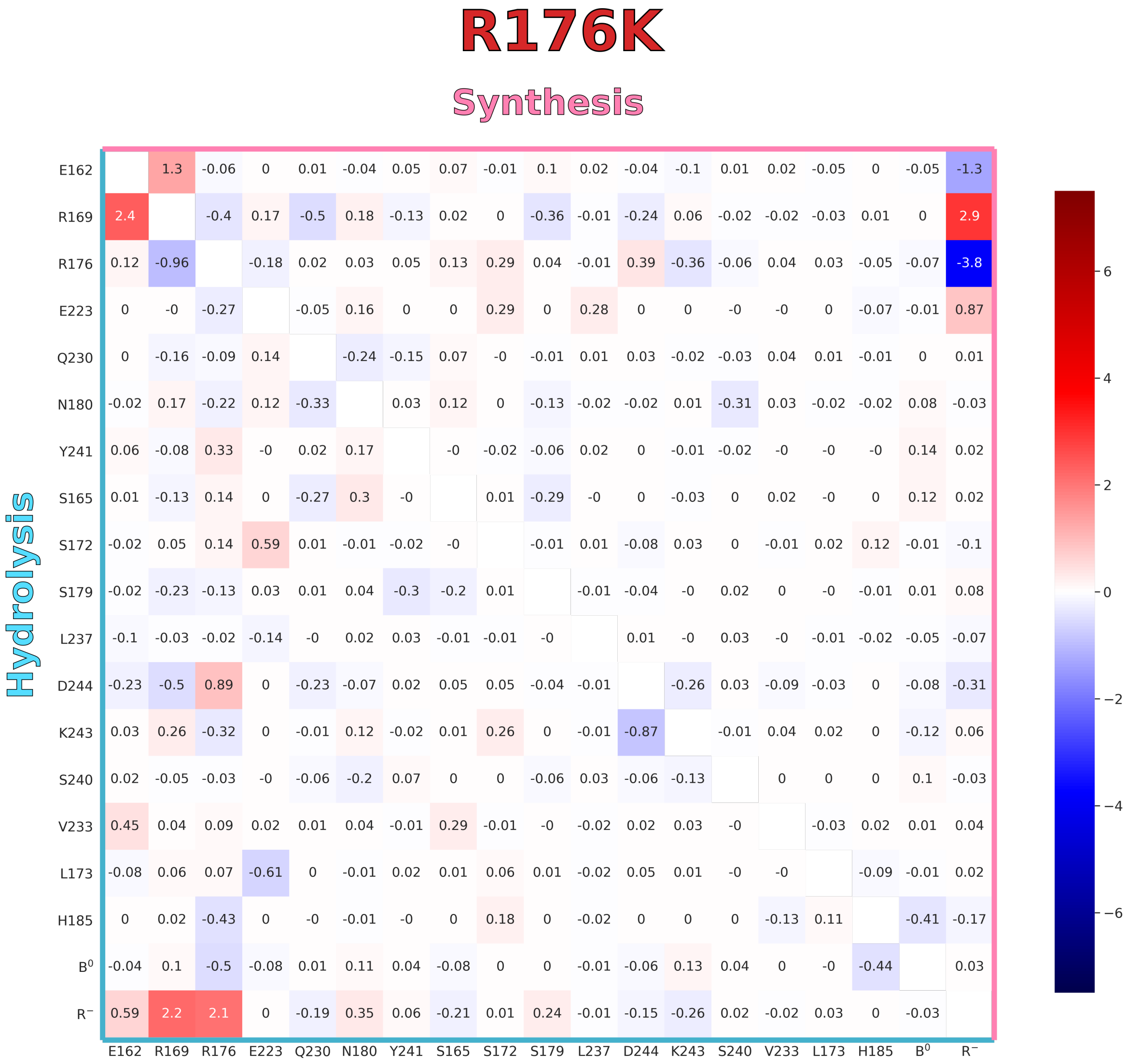


Figure S9: Average slopes (in kcal/(mol·deg)) of the selected pairwise interactions (electrostatic + vdW) as a function of the rotation angle, calculated in the 0–18° range. The values shown in the upper and lower triangle correspond to the rotation of the $F_o$ R176K mutant in the synthesis and hydrolysis direction, respectively. Locations of the residues used in the analysis are shown in Fig. S13.

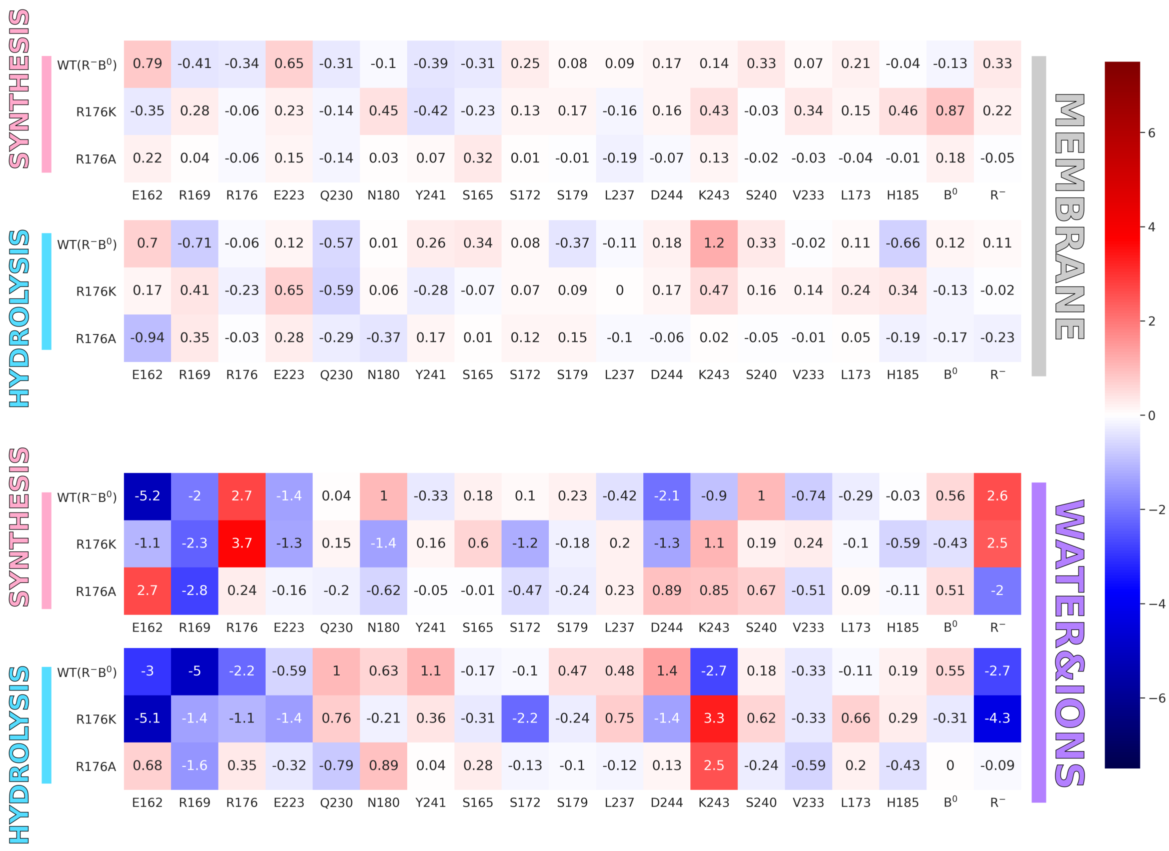


Figure S10: Average slopes (in kcal/(mol·deg)) of the sum of electrostatic and vdW interaction energies between the selected $F_o$ residues and membrane lipids (MEMBRANE) or the water solution (WATER&IONS) as a function of the rotation angle, calculated in the 0–18° range. The values corresponding to the synthesis and hydrolysis direction are labeled accordingly. Three rows in each of the matrices correspond to wild type $F_o$ ($R^{-}B^{0}$) and its R176K and R176A mutants. Locations of the residues used in the analysis are shown in Fig. S13.

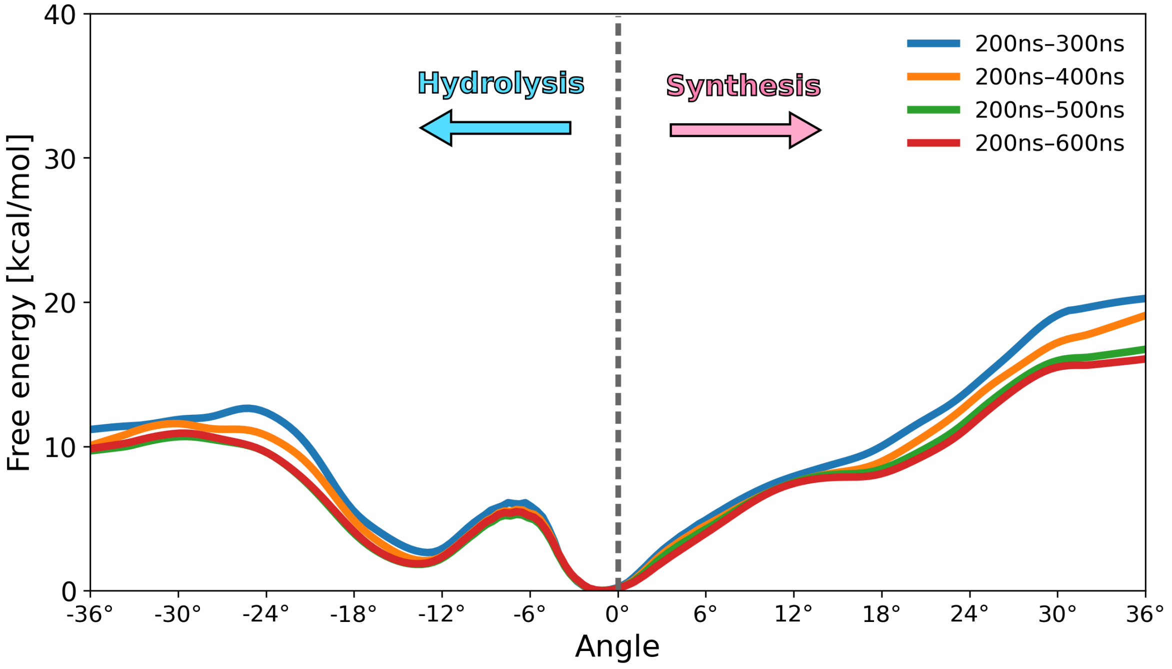


Figure S11: Convergence of the free energy profiles for the rotation of the R176A $F_o$ mutant in the synthesis and hydrolysis direction, with the length of umbrella sampling trajectories taken for analysis.

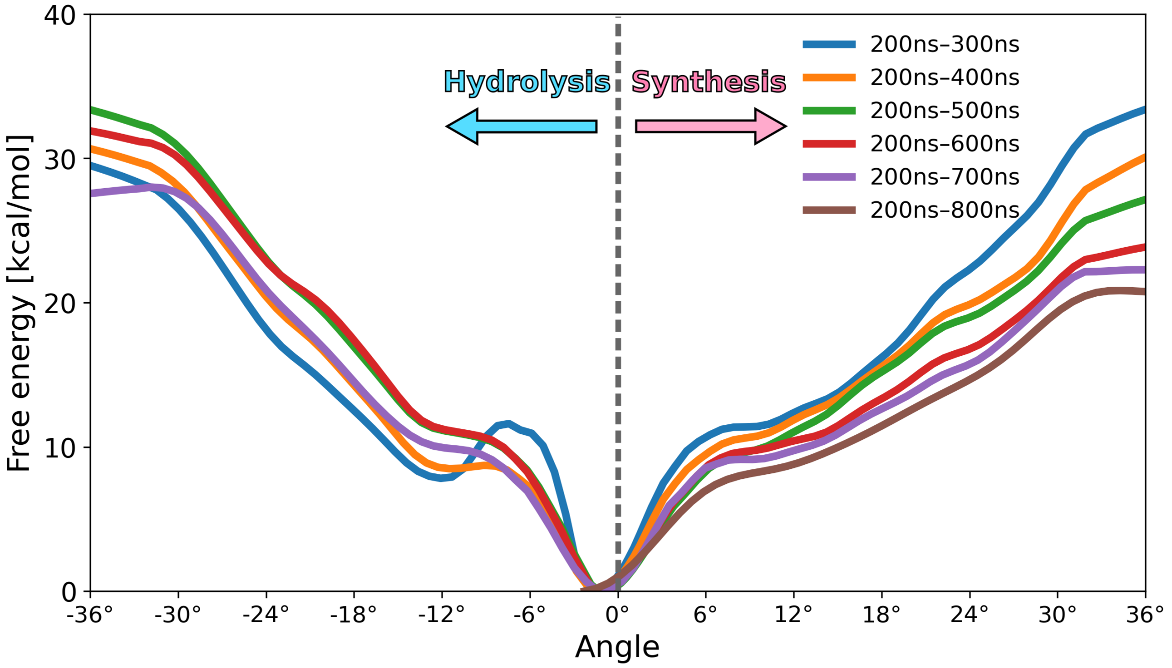


Figure S12: Convergence of the free energy profiles for the rotation of the R176K $F_o$ mutant in the synthesis and hydrolysis direction, with the length of umbrella sampling trajectories taken for analysis.

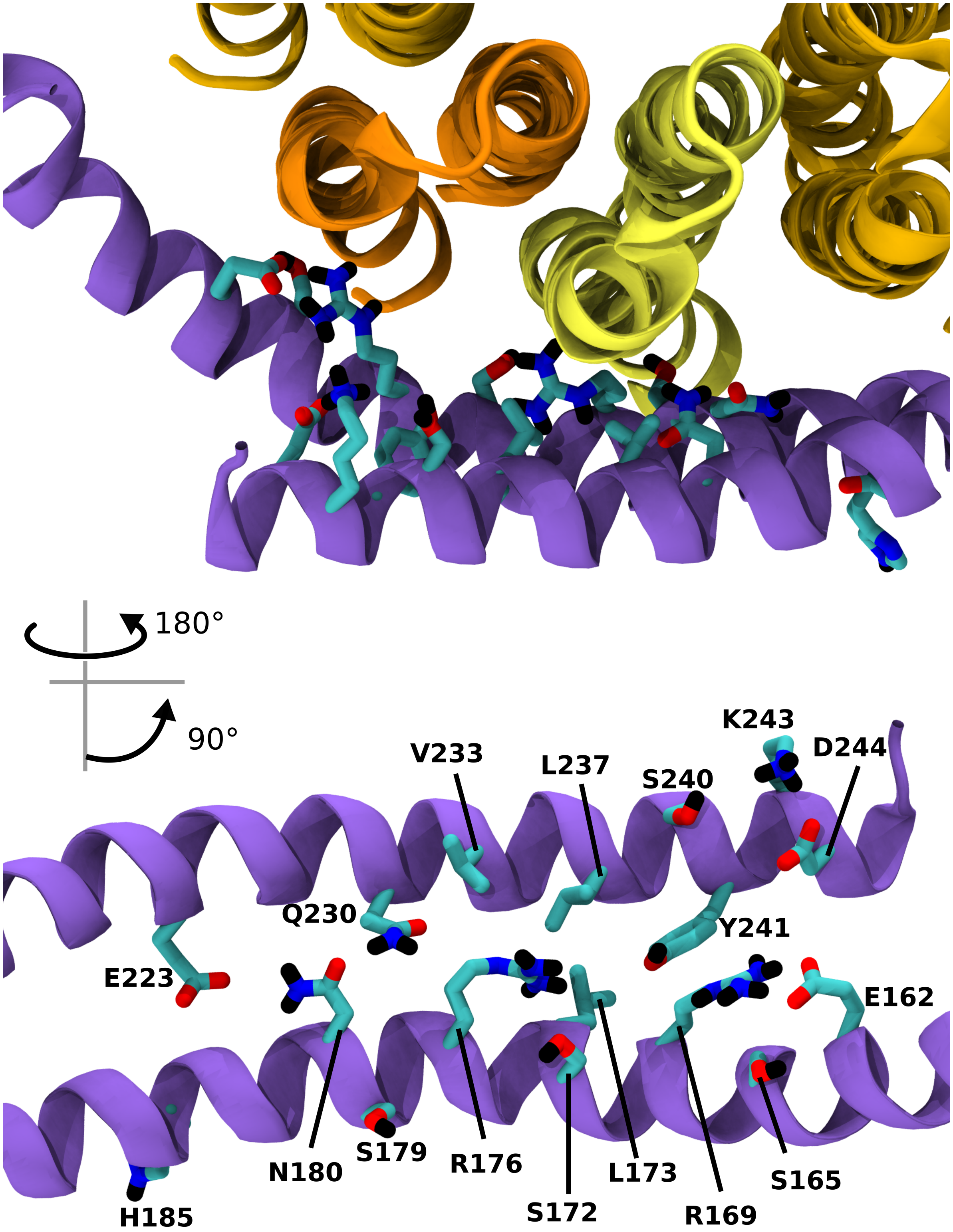


Figure S13: Residues in the a-subunit included explicitly in the analysis of pairwise enthalpic contributions to interaction free energies (Fig. S7, Fig. S8, Fig. S9 and Fig. S10). The selected residues are shown as seen from the mitochondrial matrix (top) and from the perspective of the c-ring (bottom).
